# New QTLs involved in the control of stigma position in tomato

**DOI:** 10.1101/2024.10.11.617926

**Authors:** Alessandro Riccini, Fabrizio Olivieri, Barbara Farinon, Frederique Bitton, Isidore Diouf, Yolande Carretero, Salvador Soler, Maria del Rosario Figàs, Jaime Prohens, Antonio Jose Monforte, Antonio Granell, Mathilde Causse, Andrea Mazzucato

## Abstract

The tomato mating system was strongly affected by domestication events. Mutations disrupting self-incompatibility paralleled by changes retracting the stigma position (SP) within the staminal cone conferred strict autogamy and self-fertility to the cultivated forms. Although major genes affecting these changes have been identified, SP control in domesticated forms retaining a non-inserted or a heat-inducible SP needs elucidation. To widen the possibility of identifying SP genetic determinants, we analysed the trait in four populations (two germplasm collections, a multiparental recombinant inbred and a biparental progeny) under different environmental conditions (normal and heat stressed). Overall, 36 markers significantly associated with the trait were discovered. Several co-localizations were found, both among regions firstly reported in this work and among them and previously reported positions. This supported the reliability of the analysis. Three of such regions, in the long arms of chromosome 1, 8 and 11, were validated in an independent segregating population and candidate genes in confidence intervals were identified among transcription factors and hormone, stress and cell-wall-related genes. In conclusion, the work supported the hypothesis that the SP phenotype is controlled by different key-genes in tomato, paving the way to identifying novel players and novel mechanisms involved in the regulation of herkogamy.

**Highlight:** The study of three germplasm collections in tomato allowed the identification new significant markers for the stigma position trait while confirmed several previously reported QTLs.

## Introduction

To complete the reproductive function, flowers develop specialized structures that allow the fusion of male and female gametes. Before that, pollination is needed, and this may happen by transferring the pollen grains from different plants within outcrossing species (allogamy) or from the same flower or flowers in the same plant in selfing species (autogamy). Modifications of flower morphology during domestication often determined changes in mating systems (Barret, 2010).

In tomato (*Solanum lycopersicum* L.), the flower is hermaphrodite typically with six-eight stamens closed by intermingling lateral hairs forming a cone around the pistil. The pistil is formed by two or more fused carpels and consists of a basal part, the ovary, containing the ovules with the sexual germlines, and the style, an elongated structure tipped by the expanded stigma surface. Wild tomato relatives show a stigma protruding from the staminal cone (exserted stigma) and self-incompatibility, two traits that favour allogamy (Peralta and Spooner, 2000; Li and Chetelat, 2010). Domestication events caused the “selfing syndrome”, that included the disruption of self-incompatibility and the evolution of homomorphic herkogamy, finally transforming the crop into a strict selfer (Georgiady *et al*., 2002; Chen *et al*., 2007; Boucher *et al*., 2024). Other floral changes occurred during tomato domestication, due to several human-selected mutations affecting ovary morphology, that caused an increase in fruit size and a variation in shape (van der Knaap *et al*., 2014).

The ”selfing syndrome” affected the relative length of pistil and stamens, progressively bringing the stigma position (SP) inserted within the antheridial cone (Mata-Nicolás *et al*., 2020). Although SP is a relative concept, being the result of the developmental dynamics of stamens, style and ovary, the correlation between SP and style length is generally higher than that between SP and the length of other reproductive organs (Levy *et al*., 1978; Xu *et al*., 2017). Indeed, QTLs controlling style length and SP often co-localized (Driedonks *et al*., 2018), indicating that SP is mainly determined by factors affecting style growth.

The “selfing syndrome” was driven by several changes and a polygenic control of SP has been reported (Rick, 1983; Xu *et al*., 2017). A major quantitative trait locus (QTL) was identified on the long arm of chromosome 2 and referred to as *stigma exsertion 2.1* (*se2.1*; Bernacchi and Tanksley, 1997). The gene underlying *se2.1* was later cloned and named *Style2.1* (*Solyc02g087860*); it encodes a polypeptide bearing a helix-loop-helix (HLH) motif involved in the control of cell expansion in the style (Chen *et al.,* 2007). Recently, two additional genes have been involved in SP control in tomato; *Se3.1* (*Solyc03g098070*), that encodes a C2H2 zinc finger transcription factor, emerged from the screening of a population including wild and cultivated germplasm (Shang *et al*., 2021) and *SlLst* (*Solyc12g027610*), encoding an ethylene receptor protein identified after functional characterization of a temperature sensitive mutant (Cheng *et al*., 2021).

Other SP-associated QTLs have been reported using interspecific populations. Three loci were found on chromosome 4, 8 and 9 using a cross within *S. pimpinellifolium* (*sty4.1*, *sty8.1* and *sty9.1*; Georgiady *et al*., 2002) and one on the long arm of chromosome 5 using a cross with *S. habrochaites* (*se5.1*; Gorguet *et al*., 2008). More recently, three positions controlling SP and style length were detected after exposing plants to high temperatures, which located on chromosome 1 (*qSP1*/*qSL1*), 2 (*qSL2*) and 3 (*qSP3*/*qSL3*; Xu *et al*., 2017). Notably, *qSL2* mapped to the same region of *Se2.1.* The *Se2.1* locus was also reported in the screening of recombinant inbred and introgression lines derived from a cross between *S. lycopersicum* and *S. pimpinellifolium* (Gonzalo *et al*., 2020). Although an inserted SP is a standard trait of modern tomatoes, some traditional cultivars and landraces have retained an exerted or flush stigma (Cortés-Olmos *et al*., 2015; Farinon *et al*., 2022). Studies with interspecific crosses have not yet unveiled this part of the SP trait variation.

An exserted SP is not only a constitutive feature of some tomato types, but also a heat stress related phenotype. Several studies indicated that style elongation is a genotype-dependent reaction to high temperatures, resulting in poor fertilization and low fruit set (Levy *et al*., 1978; Pan *et al*., 2017; Cheng *et al*., 2021). Because environmental conditions are important for the reproductive success in tomato and in other crop species (Ayenan *et al*., 2019), the dissection of the genetic bases of stress-related SP sensitivity represents a key-point for protecting crop yield in the context of increasing episodes of weather extremes.

Conversely, an exserted SP has been regarded as a positive trait to facilitate hybrid seed production in autogamous cereals (Marathi and Jena, 2015; Muqaddasi *et al*., 2016) and legumes (Lin *et al*., 2020). This possibility was also considered in tomato; harnessing an exerted SP combined with male sterility was proposed to ease the production of hybrid seed (Scott and George, 1980; Cheng *et al*., 2021). However, the dominance of exserted over inserted phenotypes and the incomplete penetrance of the exserted SP trait limited the use of herkogamy for producing reliable tomato hybrids (Scott and George, 1980; Rick, 1983).

Therefore, there is considerable interest in a deeper dissection of the genetic control of SP in tomato. Whereas QTL analysis was initially carried out using interspecific biparental populations, the development of high-throughput genotyping platforms and the availability of variation collections has opened the perspective for association mapping, that addresses natural genome-wide distribution of markers and alleles underlying phenotypic traits (Georgiady *et al*., 2002; Gupta *et al*., 2019). Genome-wide association studies (GWAS) have therefore become a powerful tool to study quantitative traits (Tam *et al*., 2019). GWAS relies on linkage disequilibrium (LD), the non-random co-occurrence of two or more alleles between proximal loci eventually broken down by recombination. In cultivated tomato, the extent of LD is relatively high, making it possible to perform GWAS using fewer markers than with species having lower LD (Mazzucato *et al*., 2008; Ranc *et al*., 2012; Xu *et al*., 2013). As a drawback, high LD allows a lower resolution and germplasm collections are often strongly structured and present minor frequency alleles.

Multi-parent populations, that require crosses between more than two parental lines to generate a recombinant inbred progeny, have been developed to increase the rate of LD decay. Among them, multi-parent advanced generation inter-cross (MAGIC) has the advantages of the absence of structure and the balanced allelic frequencies (Kover *et al*., 2009). Such populations can therefore increase the length of genetic maps and reduce confidence intervals compared to biparental progenies (Pascual *et al*., 2015; Arrones *et al*., 2024).

To increase the possibility of identifying SP-associated markers, we investigated SP variation in four tomato populations featuring different genetic backgrounds, encompassing the genetic diversity of wild, semi-wild and cultivated tomato. SP phenotypic values were collected under different growth conditions and used to detect QTLs by using association mapping approaches.

## Materials and methods

### Plant material

Four different tomato populations were used in the study. The Traditom Core Collection (TRA) was established in the frame of the H2020 European project “Traditional tomato varieties and cultural practices” and included 217 accessions selected to represent most of the phenotypic and genotypic variability of a larger collection (Pons *et al*., 2022). TRA was mainly composed by *S. lycopersicum* landraces, gathered from Spain, France, Italy, and Greece; moreover, three control breeding lines were included, provided by the University of Jerusalem (Pons *et al*., 2023). The *S. lycopersicum* var. *cerasiforme* population (CER) was previously constituted and characterized (Albert *et al*., 2016). In this study 132 CER accessions were used to encompass the genetic diversity of the small-fruited tomato; they included 12 *S. pimpinellifolium*, 101 *S. lycopersicum* var. *cerasiforme* and 19 admixed accessions, containing commercial cherry tomatoes and admixed genotypes. The MAGIC population (MAG) was previously developed, characterized, and genotyped (Pascual *et al*., 2015); it included 255 lines derived after intercrossing eight founder lines, four from the small-fruited group *S. lycopersicum.* var. *cerasiforme* and four from the large-fruited germplasm of cultivated tomato. Finally, the segregant interspecific population (SIP) was an F_2_ progeny (n=96) derived from the cross between *S. pimpinellifolium* (LA1589, with exserted stigma) and *S. lycopersicum* (LA1563, with inserted stigma). This population was used to validate SP-related QTLs identified by association analyses.

### Plant growth conditions

For the TRA population, four plants per accession were grown in open field in Spain (UPV, 39°28’ N 0°22’ E, ESP), in France (INRAE Avignon, 43.560’ N, - 4°51’ E, FRA) and in Italy (University of Tuscia Experimental Farm, Viterbo, 42°260′N, 12°040′E, ITA). CER was grown in France in the same conditions of TRA (Bineau *et al*., 2021), whereas MAG was grown in France in two years in greenhouse under control (MAG_N; Pascual *et al*., 2015) and high temperature conditions induced by two-months delayed sowing (MAG_H; Bineau *et al*., 2021). The SIP population, together with four plants each of the two parents and of the F1 hybrid, was grown as ITA.

In all experiments, plants were grown in plots adopting standard agronomic practices. Temperature data have been recorded during plant growth and weekly average minimum, mean and maximum values were reported.

### Plant phenotyping

In all experimental fields, except in MAG_N, SP was recorded based on scoring scale as follows :1, stigma inserted; 2, stigma at the level of the anther cone (flush stigma); 3, stigma slightly exserted (≤ 2 mm); 4, stigma highly exserted (> 2 mm; Riccini *et al*., 2021). In MAG_N, SP was recorded adopting a 1 to 3 score (1, stigma inserted; 2, flush stigma; 3, stigma exserted). In TRA, CER and SIP, SP scoring was carried out two times during the cultivation, at the beginning of flowering (from 1^st^ to 3^rd^ truss) and at the late season (from 6^th^ to 9^th^ truss). As temperatures increased during the growing season (Supplementary Fig. S1), the two SP measurements corresponded to increased heat stress experienced by the plants and were thus referred to as normal (N) and heat stress (H) conditions. Scoring was thus coded with the name of the trial followed by the growth condition abbreviation (i.e., in TRA population ESP_N, describes the SP recorded in ESP at normal conditions). As said, MAG was grown in two different years, that represented the N and H conditions (Pascual *et al*., 2015; Bineau *et al*., 2021).

To study the correlation of SP with other fertility-related traits, the number of commercial fruits collected from the 1^st^ to 4^th^ truss (FN) and the mean commercial fruit weight (FW) were retrieved from the Traditom project dataset (Pons *et al*., 2023) for 186 accessions phenotyped in Italy after removing taxa with missing data. In the same field, fruit fasciation (FASC) was scored (1, absent; 3, scarce, less than 5% of fruits affected; 5, intermediate, between 5% and 20%; 7, abundant, more than 20%). Finally, the fruit shape index (SI, ratio between polar and equatorial diameter measured on eight representative fruits) and the number of seeds per fruit (SxF, estimated after weighting all seeds extracted from a sample of 5 to 15 fruits) were calculated. All data were taken on an accession basis.

The SIP population was phenotyped on a single plant basis using the 1 to 4 SP scoring scale. Also, in this field temperatures were raising with the season (Supplementary Fig. 1F), and we could refer SP scoring to normal (SIP_N) and heat stress (SIP_H) conditions.

### Statistical analyses on phenotypic data

For TRA and CER, SP phenotypic plasticity was calculated for each accession as the difference between SP at high temperatures and SP at normal temperatures conditions, to obtain a ΔSP value for each field. For MAG, the ΔSP value (MAG_ ΔSP) was calculated by subtracting the MAG_N from the MAG_H score of each line, after converting the latter data into the 1 to 3 scale. Moreover, for TRA and CER, SP means were calculated, averaging SP under normal and heat stress conditions (ESP, ITA, FRA and CER). For SIP, SP values were mediated for each genotype in the two evaluation conditions.

The reproductive and fertility-related traits retrieved from the ITA trial were analysed for pair-wise Spearman correlations using the PROC CORR procedure of the SAS software package (SAS Institute, 1988) and correlation coefficients reported using Heatmapper (http://www.heatmapper.ca).

TRA accessions were grouped into 12 typologies according to their fruit shape, as described in Pons *et al*. (2023), with slight modifications. Typologies included genotypes with flat (big or medium according to fruit size), rectangular, ellipsoid, obovoid, round (big, medium and small), oxheart, long (San Marzano and horn) and bell pepper fruits. Since SP data did not meet the assumption of normality of residuals and no data transformation could correct this departure, a non-parametric Kruskal-Wallis test was performed using the PROC NPAR1WAY procedure with the DSCF option to compute Dwass, Steel, Critchlow-Fligner multiple comparison analysis (SAS Institute, 1988) to evaluate SP differences within each field.

### Genotypic data

TRA was genotyped using Genotyping-by-Sequencing analysis as reported (Blanca *et al*., 2022). After filtering for a minor allele frequency threshold of 5%, a maximum missing value per site and per accession of 30% and 25% respectively, and a maximum heterozygosity per site of 50%, the final dataset included 2,708 single nucleotide polymorphism (SNP) markers and 195 accessions. In addition, only biallelic loci were retained and sites with heterozygotes below 0.05% were set to missing. The tomato genome version SL2.50 was used to assign the positions of the SNPs in the analysis and markers were named reporting the number of chromosome and its position (i.e., S01_02349292 for a SNP at base pair 2,349,292 on chromosome 1).

For CER, SolCAP genotypic data were retrieved from Albert *et al*. (2016). The maximum rate of missing data was fixed at 10% and 25% per site and per accessions respectively, and a minor allele frequency threshold of 4% was applied to discard markers with rare alleles. Sites with more than 50% and less than 0.05% heterozygotes were discarded.

Genotypic data of MAG were also retrieved from a previous publication (Pascual *et al*., 2015). Briefly, 1,536 SNPs were selected from more than four million markers detected by resequencing the genomes of the eight founder lines (Causse *et al*., 2013), to construct a genetic map. SNP genotyping was performed by KASPar and Fluidigm technology as described (Pascual *et al*., 2015).

SIP genotyping was carried out by single marker analysis by PCR and restriction analysis as detailed below.

### Association mapping and linkage disequilibrium

For TRA, the analysis was carried out using 195 accessions and 12 variables, corresponding to the SP mean value, the values in N and H conditions and the ΔSP values of each field. SNP data were pruned when R^2^ was above 0.50. GWAS was performed using a mixed linear model, considering the kinship matrix and the structure obtained by Principal Component Analysis (5 components) as described (Yu *et al,* 2006). The calculation of kinship, principal component vectors and the following GWAS analysis were performed using TASSEL v. 5.2.52 (Bradbury *et al*., 2007). A Benjamini and Hochberg (1995) procedure was used to control for false discovery rate at 0.05. GWAS for CER was performed as in TRA, using four variables (CER, CER_N, CER_H and CER_ΔSP).

The 255 MAG lines were analysed using two variables (MAG_N and MAG_H). QTL mapping was carried out using the interval mapping procedure with the R package mpMap as reported by Diouf *et al*. (2018). The mpIM command was used to perform a simple interval mapping based on the regression of phenotype on the parental probabilities and estimates an allelic effect for each parent.

All significant markers were finally converted and reported according to their position in the SL4.00 version of the genome.

### QTL validation

To validate selected significant QTLs identified in the analyses, primer pairs targeting five chromosomal regions falling in QTL positions or in their proximity were designed for PCR test on the SIP population (Supplementary Table S1). DNA was extracted from the two parents, the F_1_ hybrid and 96 F_2_ progeny plants according to Fulton *et al*. (1995). PCR was carried out in 10 μL, containing 5 μL di GoTaq® Green Master Mix (Promega, Madison, WI, USA), 1 μL of each primer 10 μM and 1 μL of template DNA and using a MyCycler Thermal Cycler (BioRAD, Hercules, CA, USA) with the following program: 95°C for 2 min; 95°C for 30 s, 55-58°C depending on the primer pair for 30 s and 72°C for 20 to 60 s depending on the primer pair, looped for 30-40 times (Supplementary Table S1). For markers developed as cleaved amplified polymorphic sequences (CAPS), a final volume of 20 μL, containing 0,5 μL of the appropriate enzyme, 2 μL of buffer and 5 μL of PCR product was digested and analysed by agarose gel electrophoresis. All markers were checked for the Mendelian segregation and Spearman’s correlation was performed to estimate the significance of the validation.

Confidence intervals (CIs) were only estimated for validated markers. In detail, LD was calculated between all the unlinked loci over different chromosomes, using Plink 1.09 (Purcell *et al*., 2007); then, the 95^th^ percentile of the unlinked r^2^-distribution was regarded as the LD threshold for the following analysis. According to Breseghello and Sorrells (2006), r^2^ was plotted against base pair positions of each marker and a smooth line was drawn by second-degree loess. The number of genes within each CI was identified from the tomato genome annotation (ITAG4.0). Genes included in estimated CIs were screened for style-specificity according to expression data reported in CoNekT (https://evorepro.sbs.ntu.edu.sg/, accessed 06 June 2024, SPM value 0.70), or for flower-specificity according to TomExpress (https://tomexpress.gbfwebtools.fr, accessed 05 July 2024). Genes already reported as differentially expressed in previous analyses involving tomato pistils were evidenced (Pan *et al*., 2017; Pan *et al*., 2019; Riccini *et al*., 2021).

In addition, for the validated regions, the CIs as reported for the MAGIC marker involved were also investigated. Genes were filtered as reported (Diouf *et al*., 2018) and polymorphisms classified for the predicted high, low, moderate and modifier impact.

## Results

### Variation of stigma position in the studied tomato germplasm

The three populations studied for association mapping showed wide SP phenotypic variation. In TRA, SP ranged from 1 to 4; the accessions showing the highest SP belonged to the flat_big, oxheart, and bell pepper types (Table 1; Fig. 1; Supplementary Table S2). Similarly, high SP variation was found in CER, with two accessions of *S. pimpinellifolium*, two accessions of *S. lycopersicum* var. *cerasiforme* and one admixed accession expressing the highest value (Table 1; Supplementary Table S3). Also, the MAG population widely varied for the trait, ranging from inserted to completely exserted SP in several lines in both years of observation (Table 1; Supplementary Table S4).

**Table 1.** Mean and range of stigma position (SP) scores and accessions showing the highest values recorded in the Traditom (TRA), *S. lycopersicum* var. *cerasiforme* (CER) and MAGIC (MAG) populations grown in different locations. Accession code, name and origin are reported.

| Popula-<br>-tion | Field | No. of<br>accessions | SP score |  |  | Genotypes with highest SP scores |  |
| --- | --- | --- | --- | --- | --- | --- | --- |
|  |  |  | Mean | Min | Max | Code | Name and origin |
| TRA | ESP | 217 | 1.67 | 1 | 4 | TRBA1940 | Cor de bou, Spain |
|  |  |  |  |  |  | TRVI1830 | Bell pepper, Italy |
|  | FRA | 217 | 1.66 | 1 | 3 | TRBA1930 | Cor de bou, Spain |
|  |  |  |  |  |  | TRCA0910 | Montserrat, Spain |
|  |  |  |  |  |  | TRMO0760 | Poivron Jaune, France |
|  |  |  |  |  |  | TRVI1810 | Bell pepper, Italy |
|  |  |  |  |  |  | TRVI1830 | Bell pepper, Italy |
|  | ITA | 217 | 1.73 | 1 | 4 | TRBA1930 | Cor de bou, Spain |
|  |  |  |  |  |  | TRMO0510 | Marmande, France |
| CER |  | 136 | 1.94 | 1 | 4 | ECU1007 | *S.l. cerasiforme*, Ecuador |
|  |  |  |  |  |  | LA1245 | *S. pimpinellifolium*, Ecuador |
|  |  |  |  |  |  | LA1582 | *S. pimpinellifolium*, Peru |
|  |  |  |  |  |  | Mex-114 | *S.l. cerasiforme*, Mexico |
|  |  |  |  |  |  | Mixture | *S. pimpinellifolium*, nd |
| MAG | MAG_N | 255 | 0.42 | 1 | 3 | - | 21 lines scored 3 |
|  | MAG_H | 255 | 2.19 | 1 | 4 | - | 24 lines scored 4 |

ESP, Spain, FRA, France; ITA, Italy; MAG\_N, MAGIC population under control conditions; MAG\_H, MAGIC population under high temperatures.

**Fig. 1.**
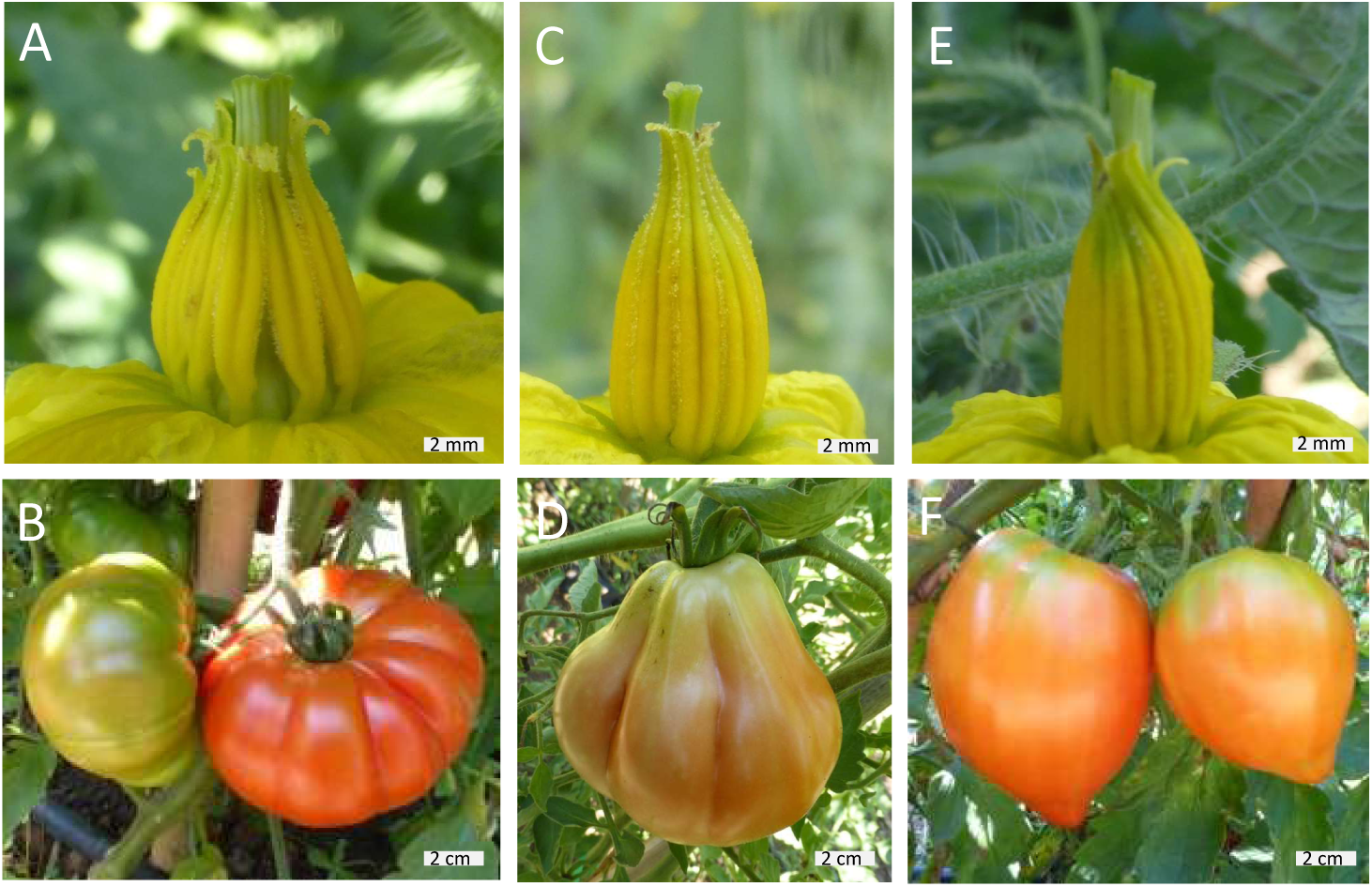
Exserted stigma phenotype and representative fruits in Traditom tomato accessions cultivated in Italy. The accessions belong to the flat_big (A, B), obovoid (C, D) and oxheart (E, F) fruit typologies. Scale bar is 2 mm in A, C, E and 2 cm in B, D, F.

The distribution of SP values differed depending on the population and on the specific trial. The frequency of accessions with inserted or flush stigma was higher in TRA field trials, with the highest percentage of inserted-SP accessions in ESP (Fig. 2A). On the contrary, the highest percentage of exserted-SP accessions was found in CER (25.8%) and MAG_H (36.5%), the two populations carrying a higher representation of the wild germplasm (Fig. 2A).

**Fig. 2.**
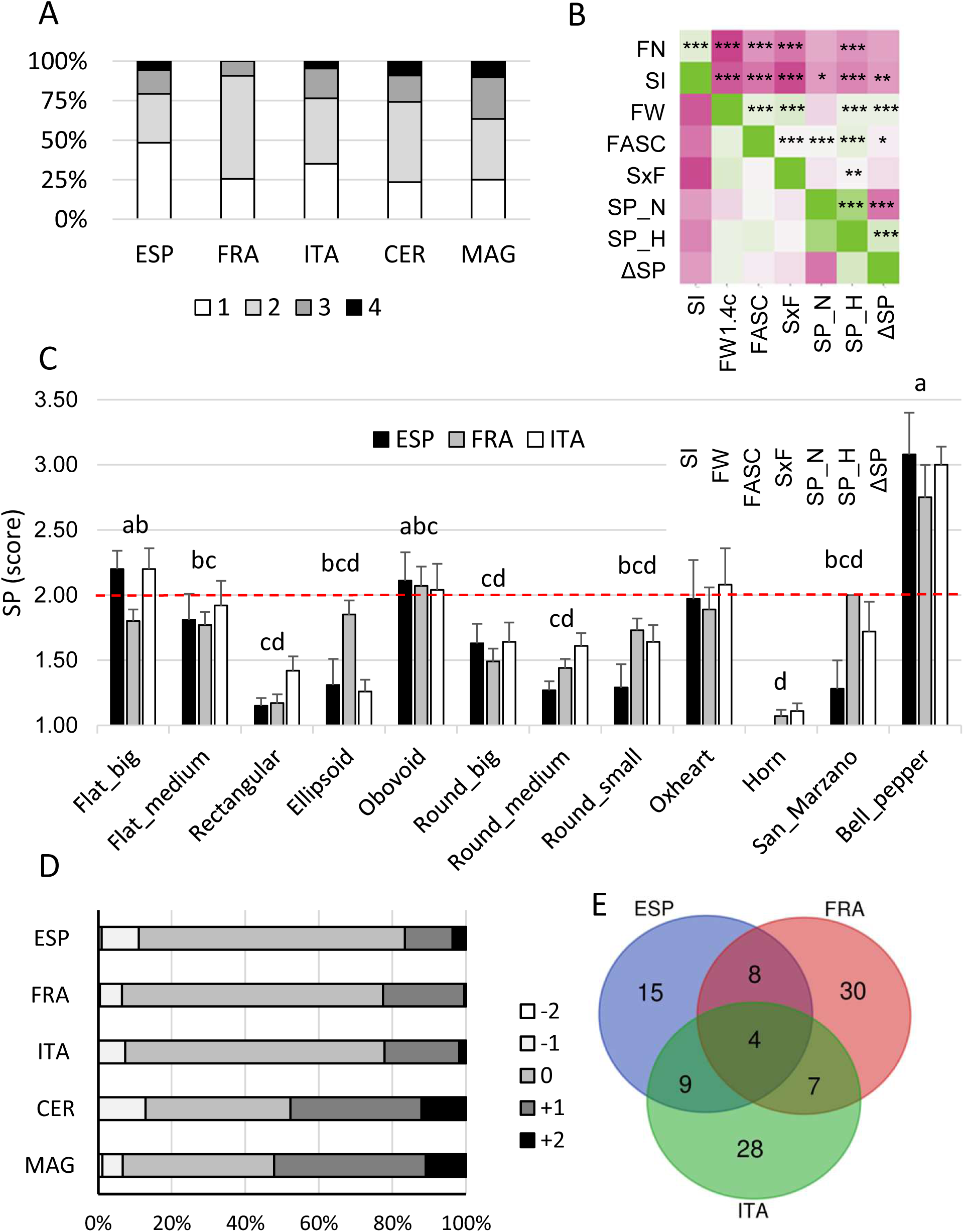
Phenotyping of stigma position (SP) in the studied populations. (A) Mean SP distribution in the Traditom (TRA) population grown in Spain (ESP), France (FRA) and Italy (ITA), in the *S. lycopersicum* var. *cerasiforme* (CER), and in MAGIC (MAG) populations. MAG represents only values from the stressed field (MAG_H). SP was scored as 1, stigma inserted; 2, stigma flush; 3, stigma slightly exserted (<2 mm); 4, stigma highly exserted (>2 mm). (B) Correlation analysis of fertility-related traits detected in the TRA collection grown in the ITA field. Magenta or green cells indicate a negative or positive correlation between traits; *, ** and *** indicate correlations with statistical significance for p < 0.05, 0.01 and 0.001 respectively. (C) SP phenotyping in the TRA collection considering the mean of two observations during the growing season in the ESP, FRA, and ITA fields, with accessions grouped in 12 typology classes. The dotted line indicates the value of the score for the flush stigma. Typologies indicated by different lowercase letters differed in all three fields after Kruskal-Wallis test and Dwass, Steel, Critchlow-Fligner multiple comparison analysis. (D) Fluctuations in SP values under increasing temperatures, shown as percentage of TRA accessions having a negative, null or positive ΔSP in the ESP, FRA and ITA trials, and in the CER and MAG populations. (E) Number of TRA accessions showing a positive ΔSP and degree of overlapping among the ESP, FRA and ITA trials.

To study the relationships between SP and other fertility-related phenotypes, data were retrieved for the TRA population grown in the ITA field (Supplementary Table S5; Pons *et al*., 2023). Strong correlations were detected among these traits and among them and SP (Fig. 2B). The SP score in stressed conditions showed the highest number of correlations, being positively correlated with FW, FASC, and SxF and negatively with SI and FN (Fig. 2B).

To describe the pattern of SP variability among different tomato fruit types, TRA accessions were grouped into 12 typologies, based on a combination of fruit shape and size (Supplementary Table S6). The non-parametric test, independently carried out for each field, always showed differences among typologies, with several significant pairwise comparisons for all three fields (Fig. 2C; Supplementary Table S6). Four typologies emerged for showing exserted SP, flat_big, obovoid, oxheart and bell pepper (Fig. 2C).

Differences between normal and stress conditions were not significant in ESP, but highly significant in the other TRA fields, with the score in stressed conditions being higher than in normal temperatures (not shown). To investigate SP stability under increasing temperatures in more detail, ΔSP was calculated. In TRA fields, most accessions were not affected by temperature variation; when fluctuations were observed, most unstable accessions showed increased exsertion under increasing temperature (Fig. 2D). Compared to TRA, the number of genotypes with SP fluctuations was higher in CER and MAG, with a proportion of genotypes showing a positive ΔSP close to or higher than 50% (Fig. 2D).

In total, 101 TRA accessions showed a positive ΔSP in at least one environment (Supplementary Table S2); 36 were reported in ESP, 49 in FRA and 48 in ITA (Fig. 2E). Four accessions (three with flat_medium and one with round_medium fruit) shared this phenotype in all environments and 24 in two environments (seven flat_big, five flat_medium, one oxheart, four obovoid, three round_big, three round_medium and one round_small). To assess the extent of within-typology SP variability, the mean values of ITA_N and ITA_H was plotted for those tomato types showing on average exserted SP. In all types, there were accessions with strongly exserted SP, at least under high temperatures, together with accessions with more inserted stigma; the bell_pepper type included only accessions with stably exserted SP (Supplementary Fig. S2).

### Genotyping and GWAS analysis in the Traditom (TRA) population

In TRA, the GWAS analysis revealed a total of 35 significant markers associated to SP, whereas no significant position was detected for ΔSP. After pooling positions in LD, 13 QTLs remained, distributed on eight chromosomes (Fig. 3; Supplementary Table S7). Three positions were detected in more than one environment. R^2^ values ranged from 8.09 to 15.43; the highest value was detected for marker S11_55195437, which was significant in two fields and three datasets (Fig. 3; Supplementary Table S7). Out of 18 QTL effects, six showed an additive and six a dominant mode of inheritance; six QTLs showed departures from both models (Supplementary Table S7). About half of the QTLs showed a higher SP in heterozygous individuals, indicating the presence of over-dominance effects (Fig. 3; Supplementary Table S7).

**Fig. 3.**
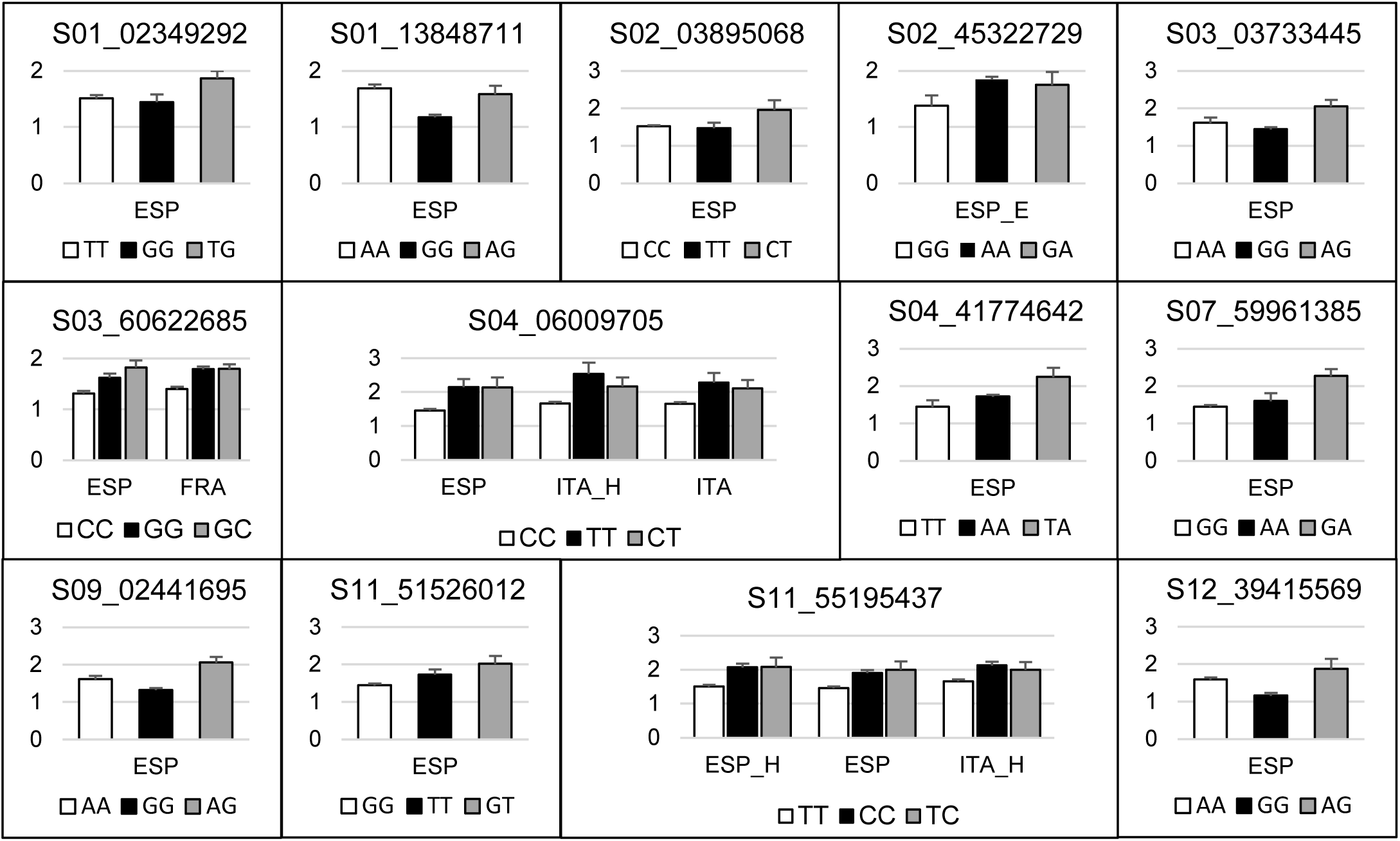
Genotypic means for the 13 QTLs identified in the Traditom core collection (TRA). TRA was analysed in three locations (Spain, ESP; France, FRA; Italy, ITA) and at two developmental times corresponding to normal (N) and heat (H) conditions. The allele found in the reference genome is reported first in all graphs (*white bar*); each bar shows the mean value + SE; y-axis represents the SP score on a 1 to 4 scale.

### Genotyping and GWAS analysis in the S. lycopersicum var. cerasiforme (CER) population

In CER, the GWAS analysis yielded 25 significant SNP loci, of which 17 identified in N and two in H growth conditions; six emerged as linked to the mean SP value. After compacting loci in LD, the positions were reduced to 11, four of which were detected in two analyses (Fig. 4; Supplementary Table S7). R^2^ ranged from 10.66 to 23.76, the highest value was detected for the QTL on the long arm of chromosome 11 (S11_55359956). Notably, all significant SNPs showed a dominant effect, although in most cases the highest score was found in heterozygous genotypes (Supplementary Table S7).

**Fig. 4.**
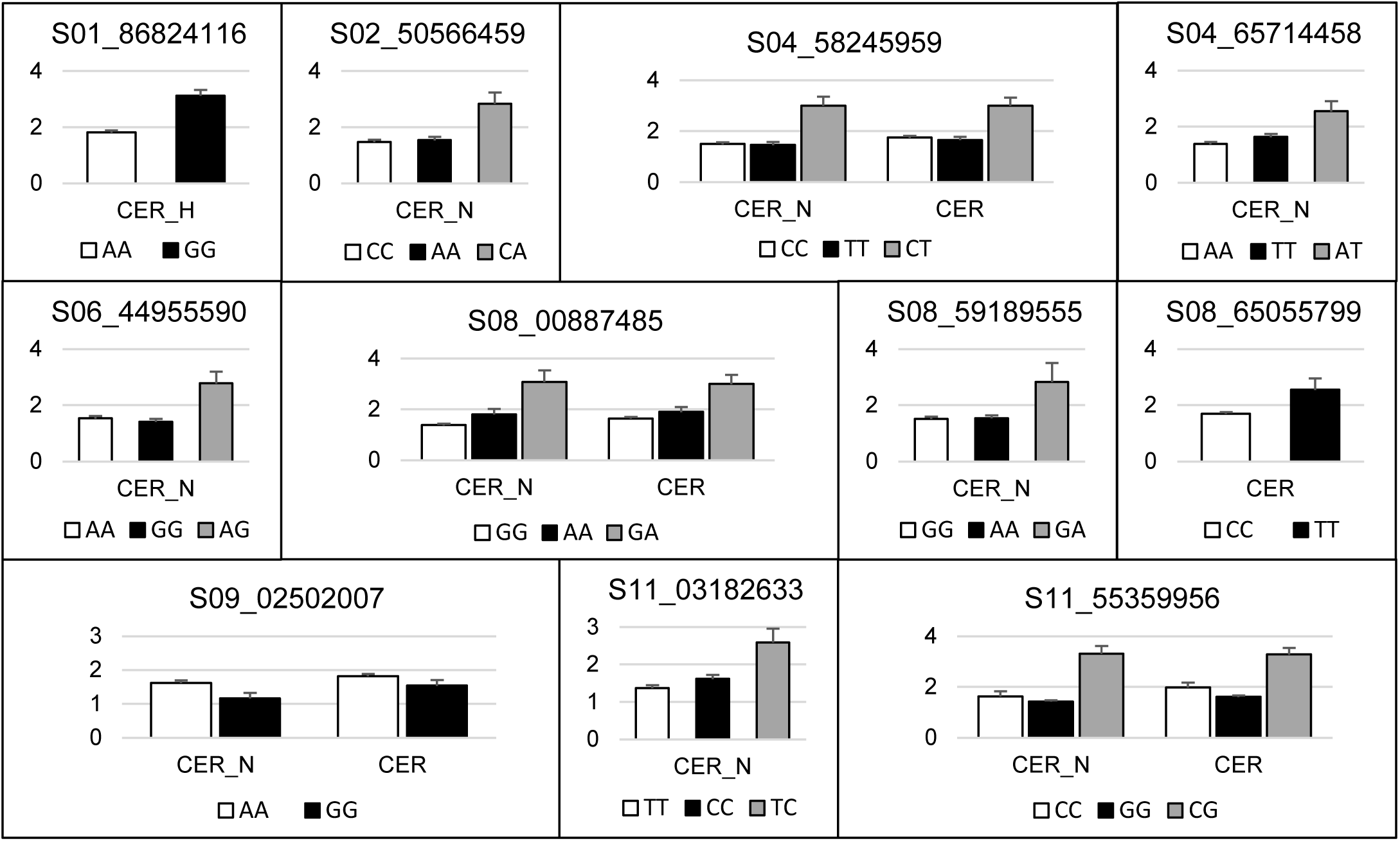
Genotypic means for the 11 QTLs identified in the *S. lycopersicum* var. *cerasiforme* (CER) population. CER was analysed in two developmental times corresponding to normal (CER_N) and heat (CER_H) temperatures. The allele found in the reference genome is reported first in all graphs (*white bar*); each bar shows the mean value + SE on a 1 to 4 scale.

### Genotyping and association analysis in the MAGIC (MAG) population

Using the datasets collected for the MAG population, 12 significant associations spread on seven chromosomes were found (Fig. 5; Supplementary Table S8). R^2^ ranged from 4.61 to 12.82, with the highest value found for X02_44814312 on chromosome 2; this, together with X07_60966290, were the only QTLs emerged in both growth conditions (Supplementary Table S8). After QTL detection, the allelic effect of the eight parental lines was estimated to see which parental alleles are in the opposite/same direction (Fig. 5; Supplementary Table S8).

**Fig. 5.**
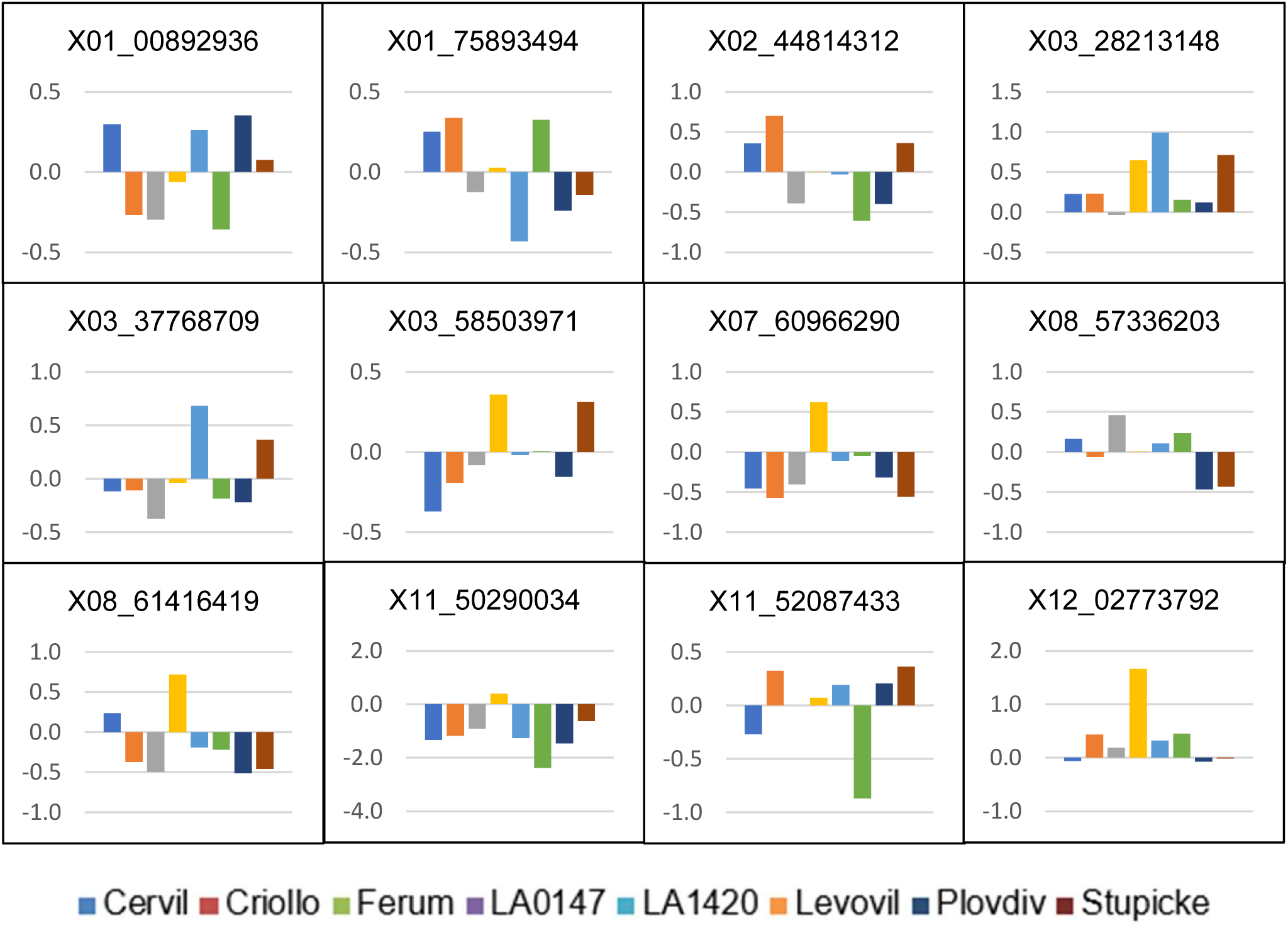
Allelic effect estimated for the 12 QTLs involved in stigma position detected in the MAGIC population. QTL effects were mean centred to facilitate the visualization of differences between parents. Marker names refer to the SL2.40 genome version.

After merging the significant positions detected in different analyses, the whole landscape of SNPs associated with SP was reported (Fig. 6). Several co-localizations were found, both among QTLs firstly reported in this work and among them and previously reported positions.

**Fig. 6.**
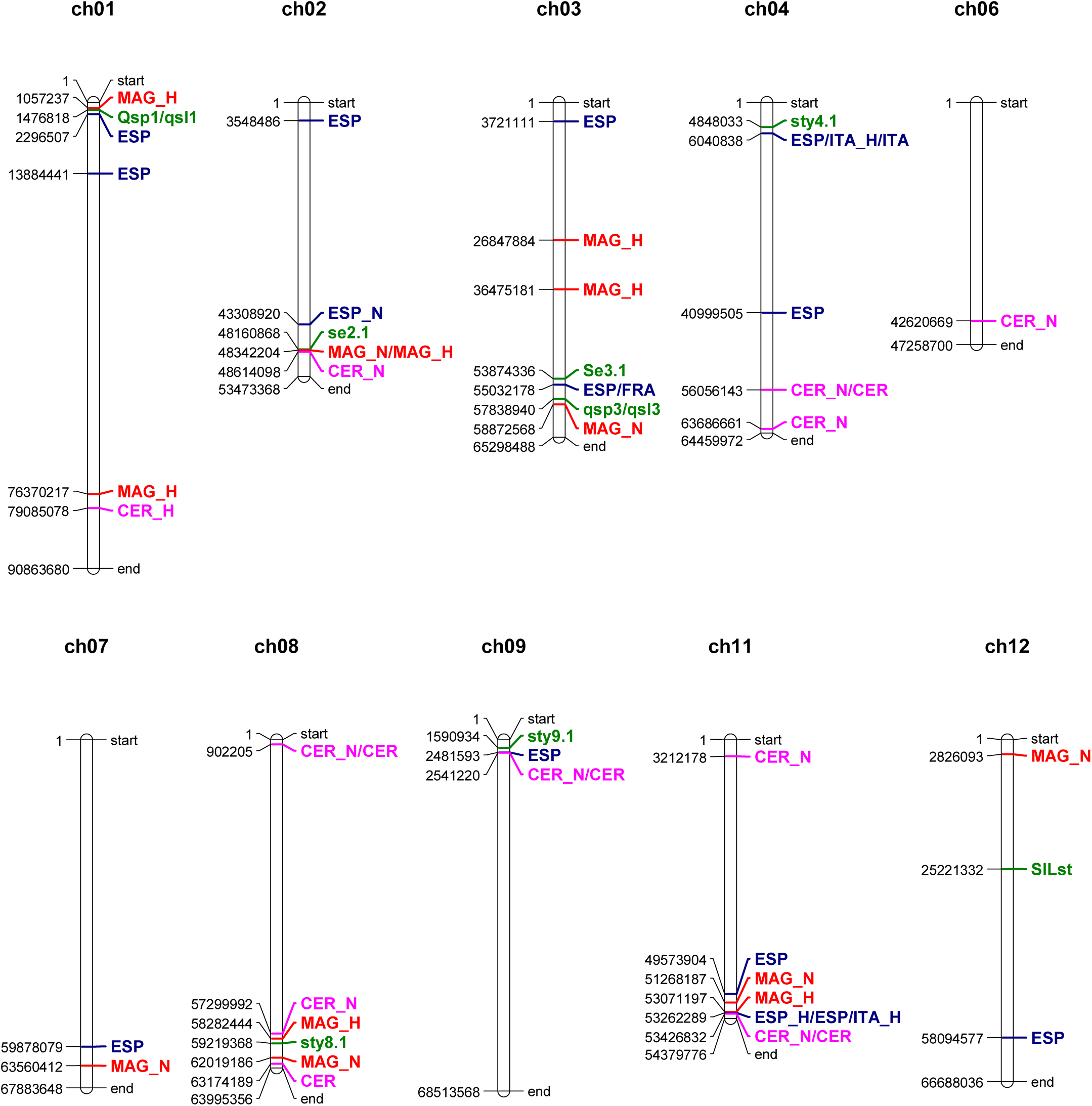
Physical map of the ten tomato chromosomes hosting significant markers associated to stigma position (SP). Markers emerged in the analysis of Traditom core collection (TRA) data scored in Spain (ESP), France (FRA) and Italy (ITA), of the *S. lycopersicum* var. *cerasiforme* (CER) collection and of the MAGIC (MAG) population. All experiments included data detected in normal (N) and heat stress (H) conditions. The position in bp (SL4.0) for each marker/gene is shown at the left side of each chromosome. Significantly associated traits are shown on the right in *blue* for TRA, *pink* for CER and *red* for MAG. Previously described QTL/genes are shown in *green*; for extended names, refer to the text.

### Validation of selected QTLs with a segregant interspecific population (SIP)

To validate GWAS results, a segregant interspecific F_2_ population (SIP) was created after crossing a *S. lycopersicum* with a *S. pimpinellifolium* genotype contrasting for SP. When phenotyped, the cultivated and wild parents stably showed the expected SP phenotype; the F_1_ had intermediate values, whereas F_2_ progeny plants showed an approximately normal segregation (Supplementary Fig. S3).

We attempted the validation of five selected QTLs, including two known (*qsp3/qsl3*, *sty8.1*) and three previously unreported positions (hereafter named *se1.1*, *se11.1*, *se12.1*; Table 2). For *se11.1* and *se12.1*, we assessed the QTL marker itself, whereas for the other positions we addressed markers in proximity or between significant QTLs. All tested markers showed Mendelian F_2_ segregation; three of them, *se1.1*, *sty8.1* and *se11.1*, resulted significantly correlated to SIP_H values. In addition, *se11.1* was also correlated to SIP_N (Table 2; Supplementary Fig. S3).

**Table 2.**
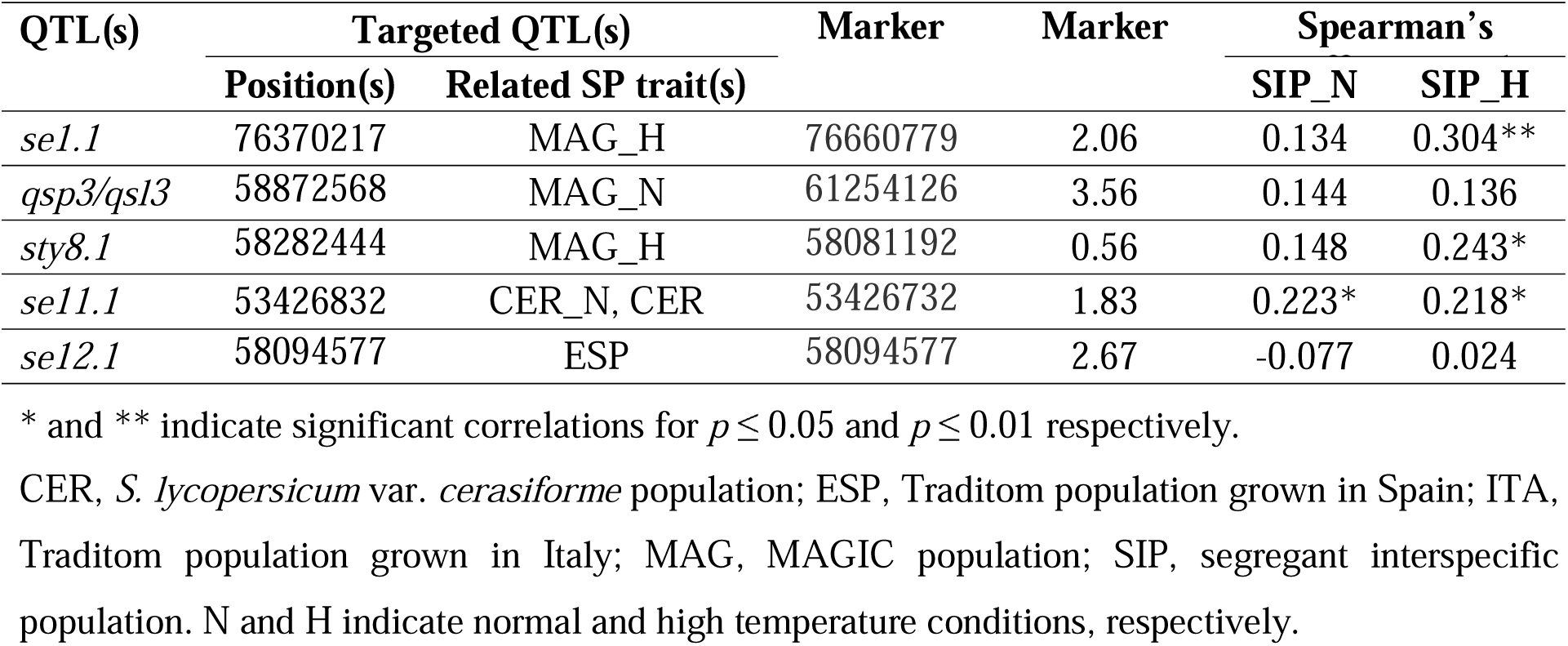
Positions of QTLs and markers chosen for validation of the association with stigma position (SP), test for Mendelian segregation and Spearman’s coefficients for the five markers in the segregant population (n=96). Marker position refers to the position of the forward primer used in PCR amplification for marker screening. All positions refer to the genome version SL4.0.

CIs calculated for the three validated markers contained 761, 661 and 221 annotated genes for *se1.1*, *sty8.1* and *se11.1* respectively (Supplementary Tables S9, S10, S11). To select candidate genes within the QTL regions we prioritized genes with reported roles in the control of pistil development, with organ-specific expression or with differential expression reported in previous works on SP regulation in tomato or in other species.

The *se1.1* CI contained five bHLH (*SlbHLH003*, *SlbHLH004*, *SlbHLH005*, *SlbHLH076* and *SlbHLH113*, according to Sun *et al*., 2015), eight MYB and 21 zinc finger transcription factors (TFs; Supplementary Table S9). In addition, we recorded four MADS-box, two homeobox, a YABBY and APETALA2 (AP2)-like TFs. Among genes involved in cell wall metabolism, we evidenced a beta-expansin precursor (*SlEXPB8*, according to Lu *et al*., 2015), linked to an expansin B15 (*SlEXPB6*). As organ differentiation and elongation is under hormone control, we highlighted two cytokinin oxidases, an auxin response factor (*ARF18*), two genes involved in gibberellin (GA) action and the calmodulin binding protein SUN3. Four DNAj proteins encoding key molecular chaperones involved in the stress response were found in this CI and, finally, seven S-adenosyl-L-methionine:salicylic acid carboxyl methyltransferase genes (S-ADEN), belonging to a gene family previously involved in plant response to abiotic stresses and in SP control (Riccini *et al.,* 2021; Supplementary Table S9).

In the *sty8.1* CI, we reported four genes encoding bHLH proteins (bHLH144, bHLH145, bHLH146 and bHLH147), three homeobox, five MYB and two NAC TFs. Also, a YABBY family member and a style-specific zinc finger protein were found. Among genes involved in abiotic stress response, we found five heat shock factors (HSF), three DNAj and two S-ADEN annotations. Of the genes involved in hormone action, we found indol-acetic acid (IAA) regulated proteins, YUCCA (*Sl-YUC4*/*SlPIN5*) and TOPLESS members, as well as genes involved in GA action, such as two GH3, a GRAS and a GA 20 oxidase 1-like (GA20ox1-like). Finally, we reported a xyloglucan endotransglucosylase/hydrolase, a xyloglucan galactosyltransferase and three expansin-like B1 genes (Supplementary Table S10).

In the *se11.1* CI, we found TFs belonging to the MADS-box, homeodomain, zinc finger, Frigida, MYB, AP2, SUN, and CLAVATA3/ESR-related gene families. Genes belonging to families involved in stress response were also reported, such as DNAj, mitogen-activated protein kinase kinase kinases, and protein kinases. As in the other CIs, we reported genes related to cytokinin, IAA and GA (GA20ox3) action. Finally, three elongation factor and an extensin-like gene were reported in this CI (Supplementary Table S11).

The *se11.1* CI spanned a region that included the *FAS* gene, whose mutation causes the increase of locule number in the tomato fruit (Xu *et al*., 2015), a phenotype positively correlated with exserted SP. To assess if the significant QTL markers in the *se11.1* position were specifically related to the *fas* mutation, we subsampled 147 TRA genotypes that showed no evidence of fasciation; the allelic effect of marker S11_55195437 remained significant in the ESP and ESP_H datasets (Supplementary Fig. S4B). Similarly, in the CER population, we subsampled 77 accessions where a wild type genotype at the *FAS* locus was reported (Blanca *et al*., 2015); in these genotypes, the allelic effect of marker S11_55359956 was also maintained, showing strong overdominance, as in the whole CER population (Supplementary Fig. 4D).

The CIs calculated on the same three regions based on the MAG markers and analysis, spanned intervals ranging from 96 to 343 kbp, containing 33, 21 and 14 filtered genes respectively (Supplementary Table S12). Only two of these genes showed variations with high predicted impact, but genes already put in evidence in the *se1.1* CI manifested variation with low, moderate or modifier impact, such as those encoding the HD-ZIP, the R2R3 MYB104 and the DNAj protein. In the *sty8.1* CI, the filtering retained the gene encoding the homeobox-leucine zipper protein ROC3. Finally, in the *se11.1* interval, we retrieved TOUSLED, a knuckle family protein and an extensin, all showing variants with possible moderate effect (Supplementary Table S12).

## Discussion

Despite its importance for the mating system and fertility, relatively few studies have analysed the control of tomato flower morphology when compared to those addressed to fruit proximal traits and primary metabolites. In the past, authors reporting QTLs controlling SP have mainly studied plant materials obtained from biparental crosses between a wild and a cultivated parent (Bernacchi and Tanksley, 1997; Georgiady *et al*., 2002; Gorguet *et al*., 2008; Xu *et al*., 2017; Gonzalo *et al*., 2020). Here, we have analysed SP associations in four populations differing in their composition and relative contribution of wild and cultivated germplasm. TRA represented the variation of European landraces, generally characterized by big fruits (87% of the accessions had a mean fruit weight > 50 g; Pons *et al*., 2023). CER sampled small-fruited tomatoes, representing spontaneous *S. lycopersicum* var. *cerasiforme*, *S. pimpinellifolium* and admixed types (98% of the accessions had mean fruit weight < 50 g; Albert *et al*., 2016). Finally, MAG included lines derived from a multiparental cross involving eight founders representing both cherry and large-fruited types.

In addition, the choice of two association analysis strategies represented a complementary approach; with GWAS we expect to detect major alleles in populations with different evolutionary history, whereas with MAGIC we exploit the variability of the founder genotypes, giving emphasis to alleles with minor frequency in the whole tomato collection. The study aimed to maximize the possibility of QTL detection and to strengthen the reliability of QTLs that were reported in more than one analysis. For validation, we adopted an interspecific population, because four out of five targeted QTLs had emerged from materials including wild germplasm.

### Stigma position showed wide variation in the studied populations

All SP classes were found in each of the studied populations. When the TRA accessions were grouped into fruit typology classes, the flat_big, obovoid, oxheart and bell_pepper genotypes showed the highest SP values. These types are characterized by multilocular and large fruits. A higher proportion of accessions with exserted SP was found in CER and MAG; in the latter, the distribution was more even, which was expected as three of the founders had exserted SP (Supplementary Table S4). Thus, accessions with exserted stigmas are found in both small-fruited wild genotypes and big-fruited cultivated typologies; the set of mutations controlling its variation is likely to only partially overlap in the two panels.

SP showed a negative correlation with FN and SI and a positive one with FW, FASC and SxF. This confirmed that in the cultivated germplasm stigma exsertion is more often found in varieties with big, fasciated, and flattened fruits, as seen in the fruit typology analysis.

### Stigma position responds to environmental variation in a genotype-specific manner

Environmental stresses affect tomato reproductive development (Sato *et al*., 2000; Giorno *et al*., 2010; Hatfield and Prueger, 2015). One of the issues of tomato growth at relatively high temperatures is the induction of stigma exsertion, that hampers self-pollination and fruit set (Saeed *et al*., 2007; Pan *et al*., 2017). Monitoring SP trends in normal and heat growth conditions revealed that SP sensitivity to high temperatures is genotype specific. In fact, we detected accessions with stable SP (inserted or exserted) and others with variable SP in different conditions.

The analysis evidenced that most temperature-sensitive accessions showed an elongation of the style under increasing temperatures, suggesting that heat stress exserts an inductive impact on style elongation. This result widens the SP-controlling candidates to genes related to abiotic stresses.

### Association analysis of stigma position in different populations revealed new significant sites while confirmed known QTLs

Three genes involved in the control of SP variation have been cloned in tomato. *Se2.1* was identified in interspecific studies and considered the major determinant in the evolution from allogamy to autogamy. Consequently, the *se2.1* mutation is essentially fixed in the cultivated tomato (Chen *et al*., 2007). Being a major gene, *Se2.1* emerged in other studies, irrespectively of the temperature regime (Xu *et al*., 2017; Gonzalo *et al*., 2000; this work). Afterwards, *Se3.1* was identified in a tomato GWAS (Shang *et al*., 2021), confirming an SP-related structural variant emerged in the analysis of bulked transcripts of genotypes with contrasting SP (Riccini *et al*., 2021; A. Riccini and A. Mazzucato, unpublished results). The *se3.1* variation was associated with the shift from flush to inserted stigma, that occurred in the domesticated germplasm (Sangh *et al*., 2021). Here, we also detected significant markers near to *Se3.1*. Finally, *SlLst*, a mutant showing exserted SP in high temperatures, was linked to an EIN4-like ethylene receptor, involving this hormone in the SP phenotype (Cheng *et al*., 2021). We did not find a position for this genetic determinant, possibly because it represents a variant which is not diffused in the tomato germplasm.

Other previously described QTLs were confirmed by our study, such as *qsp1*/*qsl1* (Xu *et al*., 2017), coincident with markers found in ESP and MAG_H, and *qSP3*, co-localized with a MAG_N marker. In addition, *sty4.1*, *sty8.1* and *sty9.1* (Georgiady *et al*., 2002) were confirmed by our analysis. Of the known QTLs or genes involved in SP control, we did not find significant markers only near *se5.1* (Gorguet *et al.,* 2008) and *SlLst* (Cheng *et al*., 2021).

Among the novel QTLs revealed by this study, the positions detected in more than one population deserve more attention. A novel QTL was found in the distal part of the long arm of chromosome 1, indicated by two markers in MAG_H and CER_H, localized within 2.7 Mbp. These positions span the overlapping region of the introgression lines IL1-1 and IL1-2, where an exserted SP has been reported (Liu and Zamir, 1999), thus supporting the existence of a QTL in the region. Five markers from eight different analyses, spanning about 4 Mbp, evidenced an unreported QTL on the long arm of chromosome 11; this position was detected in both normal and heat conditions and yielded the highest R^2^ values.

Several of the reported QTLs showed the highest mean value in heterozygotes, in accordance with reports of a high frequency of overdominance and epistatic effects in reproductive traits (Semel *et al*., 2006). More specifically, positive heterosis was reported for style length in six tomato hybrids involving a wild species in the pedigree (Nnungu and Uguru, 2014), indicating that epistatic and overdominance effects may play an important role in SP control.

### QTLs on chromosomes 1, 8 and 11 were validated in an independent biparental population

To validate GWAS results, five QTLs were tested on an interspecific F_2_ segregating SP. Two markers failed to produce a significant relationship; the one used for *qsp3/qsl3* probably because it was too far away from the QTL and *sty12.1* probably because the QTL was only found in the TRA population, which lacks the contribution of wild genomes as the validation progeny.

Differently, *se1.1* was validated in the SIP population; among the 761 genes contained in the CI, we found several TFs potentially involved in SP control, such as *SlbHLH005* (*Solyc01g090790*), that showed similarity with *SPATULA* (*SPT*) and *UNE10* (Sun *et al*., 2015), two Arabidopsis genes involved in carpel development (Heisler *et al*., 2001). This region also contained *AGL6* (*Solyc01g093960*), that encodes a protein involved in ovary development (Klap *et al*., 2016), and *AGL15*/*SlMBP11* (*Solyc01g087990*), involved in plant architecture and reproductive development, with reported effects on style growth (Guo *et al*., 2017).

Seven S-ADEN family members were detected; two of them (*Solyc01g0809903* and *Solyc01g091690*) were differentially expressed in flowers contrasting for SP (Pan *et al*., 2019; Riccini *et al*., 2021). A third member of this family, located on chromosome 9, was validated as differentially expressed in SP-contrasting genotypes (Riccini *et al*., 2021), thus supporting their role in SP control. Among the genes involved in cell wall metabolism, we found expansin B15 (*Solyc01g090820*), a pistil specific gene described in tomato fruit set (Ezura *et al*., 2017). Members of the expansin gene family have been involved in style elongation, as SPT interactors (Bernal-Gallardo *et al*., 2024). A member of the SUN gene family (SUN3, *Solyc01g088250*) was also localized in this region.

Considering the CI defined by the MAGIC marker, we found two genes showing a high impact variation, *Solyc01g090490* and *Solyc01g090500* (Supplementary Table S12). These genes encode a transcription termination factor and a lipid transfer protein. The HD-ZIP *Solyc01g090460* and the R2R3 *MYB104 Solyc01g090530*, that were mentioned as miRNA targets in heat-stressed pistils (Pan *et al*., 2017), also felt in this CI.

The marker tested on chromosome 8 was also validated; it was about 1 Mb from *sty8.1*, a QTL previously reported with a very high R^2^ value (Georgiady *et al*., 2002). The strong overdominance reported for this locus in the CER population supports the hypothesis that other factors apparently modify the effect of the QTL on the phenotype (Georgiady *et al*., 2002). Among the four bHLH members found in *sty8.1*, we focus on *bHLH147*, which is ortholog to *MYC2*, that encodes a protein involved in the abscisic and jasmonic acid pathways in Arabidopsis (Sun *et al*., 2015). Further, we found the homeobox *Solyc08g078300*, ortholog of *ARABIDOPSIS THALIANA HOMEOBOX PROTEIN 2* (*ATHB2*/*HAT4*). There is evidence that *HAT* genes are involved in the shaping of the gynoecium, being positively regulated by SPT, and interacting with indol-acetic acid (IAA)-response members (Turchi *et al*., 2015). Also, in this CI were found a CRABS CLAW family member (*Solyc08g079100*) and a style specific zinc finger protein expressed before flowering (*Solyc08g078590*). This latter gene showed co-expression with *Se2.1* (CoNekT database). In the *sty8.1* CI, the homeobox-leucine zipper protein ROC3 was retained by filtering, showing a variant with moderate impact; its paralog in chromosome 4 was reported as a pistil-specific gene (Ezura *et al*., 2017).

We reported in this region also genes involved in abiotic stresses, such as five HSF and HSF-related members that mediate plant response to heat (Kotak *et al*., 2007; Jung *et al*., 2012). These are candidate as SP-control genes, together with genes involved in the metabolic pathway of abscisic and salicylic acid, ethylene, oxidative burst, and calmodulin, that are known to play significant roles in the heat stress response (Larkindale *et al*., 2005; Zhang *et al*., 2009; Firon *et al*., 2012). Notably, it has been concluded that herkogamy can rapidly evolve in response to environmental changes, such as variation in pollinator communities or abiotic factors (Opedal *et al*., 2022).

Among hormone-related genes, we found *Sl-YUC4*/*SlPIN5*/*SlFZY2* (*Solyc08g068160*), which is ortholog to *AtYUC2* and possibly MYB-regulated (Meng *et al*., 2023), and *TOPLESS2* (*Solyc08g076030*), member of a family involved in IAA response and controlling ovary development (He *et al*., 2021). Also, genes involved in xyloglucan metabolism (*Solyc08g076080*; *Solyc08g079040*) and in cell expansion (*Solyc08g077330*, *Solyc08g077900*, *Solyc08g077910*) were reported. Such protein families have been involved in SP control (Pan *et al*., 2019; Riccini *et al*., 2021) and in early ovary development (Carrera *et al*., 2012).

Finally, the QTL found on chromosome 11 was positively validated, with a significant effect in both normal and stress conditions. The calculated CI spanned 221 coding sequences, including *SlMADS55* (*Solyc11g069770*; Hileman *et al*., 2006), a gene co-expressed with *Se2.1* (CoNekT database). *SlMADS55* showed similarity to *AGL62*, involved in the fruit set-related activation of IAA synthesis in the Arabidopsis pistil (L. Guo *et al*., 2022).

Genes possibly related to stress response were reported, such as *HSP40* (*Solyc11g071830*), that has been involved in the response to heat and cold, and to the action of plant hormones. This protein is pistil and stamen specific, with a regulation during the reproductive phases (Li *et al*., 2012); its role in heat stress response and the specific expression makes it a good candidate for controlling SP in tomato.

Among genes involved in cell wall remodelling, the region contained two elongation factors, differentially expressed between SP-contrasting genotypes (*Solyc11g072190*, Riccini *et al*., 2021; *Solyc11g069700*, Pan *et al*., 2019), a style-specific glucan endo-1,3-beta-glucosidase (*Solyc11g072230*) and SUN31 (*Solyc11g071840*), a calmodulin binding protein expressed during flower and fruit development (Wu *et al*., 2011; Sacco *et al*., 2015). A recently annotated extensin-like gene (*Solyc11g161770*) was also located in this CI. As for the other CIs, we reported genes related to hormone action, such as *ARF10A*, whose ortholog was described as a strong candidate for controlling SP in rice (N. Guo *et al*., 2022), and GA (GA20ox3, *Solyc11g072310*) and IAA (*YABBY2b*, *Solyc11g071810*) biosynthetic genes. *YABBY2b* is involved in the inversion that, including also *CLAVATA3*/*ESR-related* (*Solyc11g071380*), is responsible for the *fas* phenotype (Cong *et al*., 2008; Zhang *et al*., 2024). *FAS* controls locule number in tomato; when mutated, it confers fruit fasciation and, generally, flat shape (Rodriguez *et al*., 2011). Accessions with fasciated are frequent in TRA, and are associated with SP, because exsertion was more frequent in such fruit types. The involvement of YABBY TFs in *Mimulus lewisii* style elongation (Ding *et al*., 2021) supports a direct control of FAS on SP. However, the mechanisms underlying the correlation between YABBY TFs and style exsertion remains to be elucidated, because loss-of-function mutants showed a shorter style in *Mimulus* (Ding *et al*., 2021), but a longer style in tomato (Cong *et al*., 2008). The analysis of subsamples of individuals not carrying the *fas* mutation in TRA and CER confirmed a significant QTL on the long arm of chromosome 11, indicating that at least a second gene is involved in SP regulation in this region.

In conclusion, this work demonstrated that the SP phenotype is controlled by different key-genes in tomato. The loss of exsertion occurred with domestication through recessive mutations (i.e., *se2.1*), was later associated with genetic changes that increased stigma insertion in the modern varieties (i.e., *se3.1*). In addition, single mutations, directly or indirectly affecting style elongation, and genes responsible for stress sensitivity are likely to contribute to the phenotype in a genotype-specific manner (i.e., *SlLst*). This study indicated both known and novel loci that could be involved in these pathways. Future research will disclose the underlying genes as well as the mechanisms involved in herkogamy regulation.

## Supplementary data

The following supplementary data are available at JXB online.

**Table S1.** Markers used for the validation of QTLs associated to stigma position (SP).

**Table S2.** Phenotypic data collected for stigma position (SP) in the Traditom core collection.

**Table S3.** Phenotypic data collected for stigma position (SP) in the *S. lycopersicum* var. *cerasiforme* collection.

**Table S4.** Phenotypic data collected for stigma position (SP) in parents and lines of the MAGIC population.

**Table S5.** Fruit phenotypic data collected in the Traditom core collection grown in Italy.

**Table S6.** Classification of the Traditom core collection into fruit typology classes.

**Table S7.** SNPs significantly associated with stigma position (SP) in the Traditom core collection (TRA) and in the *S. lycopersicum* var. *cerasiforme* (CER) populations.

**Table S8.** Significant SNPs associated with stigma position (SP) in the MAGIC population.

**Table S9.** Genes annotated within the *se1.1* confidence interval.

**Table S10.** Genes annotated within the *sty8.1* confidence interval.

**Table S11.** Genes annotated within the *se11.1* confidence interval.

**Table S12.** Genes included in confidence intervals calculated for the MAGIC markers for the three validated regions.

**Fig. S1.** Temperatures registered in the experimental fields.

**Fig. S2.** Stigma position (SP) registered within selected typologies of the Traditom core collection.

**Fig. S3.** Validation of QTLs associated to stigma position (SP).

**Fig. S4.** Genotypic means for chromosome 11 SNPs associated to stigma position (SP) in whole collections and in filtered subsamples wild-type at the *FAS* gene.

## Acknowledgements

We thank the C.M. Rick TGRC, University of California, Davis, USA for seed stocks supply, Fabrizio de Angelis and Federico Pascarella for technical assistance in QTL validation, and Sotirios Fragkostefanakis for helpful discussion.

## Author contributions

AM, AG, JP, AJM, and MC: conceptualization and coordination; AG: coordination of the Traditom and Harnesstom projects; AR, ID, YC, SS, MRF: growth and phenotyping the populations; AM, AR, FO, BF, SS: curating phenotypic data; AM, AR, FO, BF, FB: performing GWAS and bioinformatic analyses; BF and FO: performing validation analysis; AR, FO, BF, AM, MC: preparing all figures, and drafting the manuscript.

## Data availability

The authors confirm that all data from this study is available in the manuscript its Supplementary Information.

## Abbreviations

ARF: auxin response factor
CAPS: cleaved amplified polymorphic sequence
CER: *S. lycopersicum* var. *cerasiforme* population
CI: confidence interval
ESP: experimental trail carried out in Spain
*FAS*: *Fasciated* gene
FASC: fasciation
FN: fruit number
FRA: experimental trial carried out in France
FW: fruit weight
GA: gibberellin
GWAS: genome-wide association study
H: high temperature growth conditions
HLH: helix-loop-helix
HSF: heat shock factor
IAA: indol-acetic acid
ITA: experimental trial carried out in Italy
LD: linkage disequilibrium
MAG: MAGIC population
N: normal growth conditions
QTL: quantitative trait locus
S-ADEN: S-adenosyl-L-methionine:salicylic acid carboxyl methyltransferase gene
SI: shape index
SIP: segregant interspecific population
SNP: single nucleotide polymorphism
SP: stigma position
*SPT*: *Spatula*
SxF: seeds per fruit
TF: transcription factor
TRA: Traditom core collection.

## References

1. Albert E, Segura V, Gricourt J, Bonnefoi J, Derivot L, Causse M. 2016. Association mapping reveals the genetic architecture of tomato response to water deficit: focus on major fruit quality traits. Journal of Experimental Botany 67, 6413–6430. 10.1093/jxb/erw411.

2. Arrones A, Antar O, Pereira-Dias L, et al. 2024. A novel tomato inter-specific (*Solanum lycopersicum* var. *cerasiforme* and *S. pimpinellifolium*) MAGIC population facilitates trait association and candidate gene discovery in untapped exotic germplasm. Horticulture Research 11, uhae154. 10.1101/2024.02.28.582481.

3. Ayenan MAT, Danquah A, Hanson P, Ampomah-Dwamena C, Sodedji FAK, Asante IK, Danquah EY. 2019. Accelerating Breeding for Heat Tolerance in Tomato (*Solanum lycopersicum* L.): An Integrated Approach. Agronomy 9, 720. 10.3390/agronomy9110720.

4. Bai Y, Lindhout P. 2007. Domestication and breeding of tomatoes: what have we gained and what can we gain in the future? Annals of Botany 100, 1085–1094.

5. Barrett SC. 2010. Understanding plant reproductive diversity. Philosophical Transactions of the Royal Society B: Biological Sciences 365, 99–109.

6. Benjamini Y, Hochberg Y. 1995. Controlling the False Discovery Rate: A Practical and Powerful Approach to Multiple Testing, Journal of the Royal Statistical Society: Series B (Methodological) 57, 289–300. 10.1111/j.2517-6161.1995.tb02031.x

7. Bernacchi D, Tanksley SD. 1997. An interspecific backcross of *Lycopersicon esculentum* × *L. hirsutum*: linkage analysis and a QTL study of sexual compatibility factors and floral traits. Genetics 147, 861–877.

8. Bernal-Gallardo JJ, González-Aguilera KL, de Folter S. 2024. *EXPANSIN15* is involved in flower and fruit development in Arabidopsis. Plant Reproduction 37, 259–270. 10.1007/s00497-023-00493-4.

9. Blanca J, Montero-Pau J, Sauvage C, Bauchet G, Illa E, Díez MJ, Francis D, Causse M, van der Knaap E, Cañizares J. 2015. Genomic variation in tomato, from wild ancestors to contemporary breeding accessions. BMC Genomics 16, 257. doi: 10.1186/s12864-015-1444-1.

10. Blanca J, Pons C, Montero-Pau J, et al. 2022. European traditional tomatoes galore: a result of farmers’ selection of a few diversity-rich loci. Journal of Experimental Botany 73, 3431–3445. doi: 10.1093/jxb/erac072.

11. Boucher JJ, Ireland HS, Wang R, David KM, Schaffer RJ. 2024. The genetic control of herkogamy. Functional Plant Biology 51, FP23315. doi: 10.1071/FP23315. PMID: 38687848.

12. Bineau E, Diouf I, Carretero Y, Duboscq R, Bitton F, Djari A, Zouine M, Causse M. 2021. Genetic diversity of tomato response to heat stress at the QTL and transcriptome levels. Plant Journal 107, 1213–1227. doi: 10.1111/tpj.15379.

13. Bradbury PJ, Zhang Z, Kroon DE, Casstevens TM, Ramdoss Y, Buckler ES. 2007. TASSEL: software for association mapping of complex traits in diverse samples. Bioinformatics 23, 2633–5. doi: 10.1093/bioinformatics/btm308.

14. Breseghello F, Sorrells ME. 2006. Association mapping of kernel size and milling quality in wheat (*Triticum aestivum* L.) cultivars. Genetics 172, 1165–77. doi: 10.1534/genetics.105.044586.

15. Carrera E, Ruiz-Rivero O, Pereira Peres LE, Atares A, Garcia-Martinez JL. 2012. Characterization of the *procera* tomato mutant shows novel functions of the SlDELLA protein in the control of flower morphology, cell division and expansion, and the auxin-signaling pathway during fruit-set and development. Plant Physiology 160, 1581–1596. 10.1104/pp.112.204552.

16. Causse M, Desplat N, Pascual L, et al. 2013. Whole genome resequencing in tomato reveals variation associated with introgression and breeding events. BMC Genomics 14, 791. doi: 10.1186/1471-2164-14-791.

17. Chen KY, Cong B, Wing R, Vrebalov J, Tanksley SD. 2007. Changes in regulation of a transcription factor lead to autogamy in cultivated tomatoes. Science 318, 643–5. doi: 10.1126/science.1148428.

18. Cheng MZ, Gong C, Zhang B, et al. 2021. Morphological and anatomical characteristics of exserted stigma sterility and the location and function of SlLst (*Solanum lycopersicum Long styles*) gene in tomato. Theoretical and Applied Genetics 134, 505–518. doi: 10.1007/s00122-020-03710-0.

19. Cleveland WS. 1979. Robust locally weighted regression and smoothing scatterplots. Journal of the American Statistical Association 74, 829–836.

20. Cong B, Barrero LS, Tanksley SD. 2008. Regulatory change in YABBY-like transcription factor led to evolution of extreme fruit size during tomato domestication. Nature Genetics 40, 800.

21. Cortés-Olmos C, Valcárcel JV, Roselló J, Díez M, Cebolla-Cornejo J. 2015. Traditional eastern Spanish varieties of tomato. Scientia Agricola 5, 420–431. 10.1590/0103-9016-2014-0322.

22. Ding B, Li J, Gurung V, Lin Q, Sun X, Yuan Y-W. 2021. The leaf polarity factors SGS3 and YABBYs regulate style elongation through auxin signaling in *Mimulus lewisii*. New Phytologist 232, 2191–2206. 10.1111/nph.17702.

23. Diouf IA, Derivot L, Bitton F, Pascual L, Causse M. 2018. Water deficit and salinity stress reveal many specific QTL for plant growth and fruit quality traits in tomato. Frontiers in Plant Science 9. 10.3389/fpls.2018.00279.

24. Doganlar S, Frary A, Tanksley SD. 2000. The genetic basis of seed-weight variation: tomato as a model system. Theoretical and Applied Genetics 100, 1267–1273.

25. Driedonks N, Wolters-Arts M, Huber H, de Boer G-J, Vriezen W, Mariani C, Rieu I. 2018. Exploring the natural variation for reproductive thermotolerance in wild tomato species. Euphytica 214, 67. 10.1007/s10681-018-2150-2

26. Ezura K, Ji-Seong K, Mori K, Suzuki Y, Kuhara S, Ariizumi T, Ezura H. 2017. Genome-wide identification of pistil-specific genes expressed during fruit set initiation in tomato (*Solanum lycopersicum*). PLoS One 12, e0180003. 10.1371/journal.pone.0180003.

27. Farinon B, Picarella ME, Mazzucato A. 2022 Dynamics of fertility-related traits in tomato landraces under mild and severe seat stress. Plants 11, 881. 10.3390/plants11070881.

28. Firon N, Pressman E, Meir S, Khoury R, Altahan L. 2012. Ethylene is involved in maintaining tomato (*Solanum lycopersicum*) pollen quality under heat-stress conditions. AoB Plants pls024. doi: 10.1093/aobpla/pls024.

29. Frary A, Nesbitt TC, Grandillo S, Knaap E, Cong B, Liu J, Meller J, Elber R, Alpert KB, Tanksley SD. 2000. *fw2.2*: a quantitative trait locus key to the evolution of tomato fruit size. Science 289, 85–88. doi: 10.1126/science.289.5476.85.

30. Fulton TM, Chunwongse J, Tanksley SD. 1995. Microprep protocol for extraction of DNA from tomato and other herbaceous plants. Plant Molecular Biology Reporter 13, 207–209. 10.1007/BF02670897.

31. Georgiady MS, Whitkus RW, Lord EM. 2002. Genetic analysis of traits distinguishing outcrossing and self-pollinating forms of currant tomato, *Lycopersicon pimpinellifolium* (Jusl.) Mill. Genetics 161, 333–344.

32. Giorno F, Wolters-Arts M, Grillo S, Scharf KD, Vriezen WH, Mariani C. 2010. Developmental and heat stress-regulated expression of *HsfA2* and small heat shock proteins in tomato anthers. Journal of Experimental Botany 61, 453–462. doi: 10.1093/jxb/erp316.

33. Gonzalo MJ, Li YC, Chen KY, Gil D, Montoro T, Nájera I, Baixauli C, Granell A, Monforte AJ. 2020. Genetic Control of Reproductive Traits in Tomatoes Under High Temperature. Frontiers in Plant Science 11, 326. doi: 10.3389/fpls.2020.00326.

34. Gorguet B, Eggink PM, Ocaña J, Tiwari A, Schipper D, Finkers R, Visser RG, van Heusden AW. 2008. Mapping and characterization of novel parthenocarpy QTLs in tomato. Theoretical and Applied Genetics 116, 755–67. doi: 10.1007/s00122-007-0708-9.

35. Guo X, Chen G, Naeem M, Yu X, Tang B, Li A, Hu Z. 2017. The MADS-box gene *SlMBP11* regulates plant architecture and affects reproductive development in tomato plants. Plant Science 258, 90–101. 10.1016/j.plantsci.2017.02.005.

36. Guo L, Luo X, Li M, Joldersma D, Plunkert M, Liu Z. 2022. Mechanism of fertilization-induced auxin synthesis in the endosperm for seed and fruit development. Nature Communications 13, 3985. doi: 10.1038/s41467-022-31656-y.

37. Guo N, Wang Y, Chen W, et al. 2022. Fine mapping and target gene identification of qSE4, a QTL for stigma exsertion rate in rice (*Oryza sativa* L.). Frontiers in Plant Science 13. 10.3389/fpls.2022.959859.

38. Gupta PK, Kulwal PL, Jaiswal V. 2019. Association mapping in plants in the post-GWAS genomics era. Advances in Genetics 104, 75–154.

39. Hatfield JL, Prueger JH. 2015. Temperature extremes: Effect on plant growth and development. Weather and Climate Extremes 10, 4–10.

40. He M, Song S, Zhu X, et al. 2021. *SlTPL1* silencing induces facultative parthenocarpy in tomato. Frontiers in Plant Science 12. 672232. doi: 10.3389/fpls.2021.672232.

41. Heisler MG, Atkinson A, Bylstra YH, Walsh R, Smyth DR. 2001. *SPATULA*, a gene that controls development of carpel margin tissues in Arabidopsis, encodes a bHLH protein. Development 128, 1089–98. doi: 10.1242/dev.128.7.1089.

42. Hileman LC, Sundstrom JF, Litt A, Chen M, Shumba T, Irish VF. 2006. Molecular and Phylogenetic Analyses of the MADS-Box Gene Family in Tomato. Molecular Biology and Evolution 23, 2245–2258. 10.1093/molbev/msl095.

43. Huang Z, Houten J, Gonzalez-Escobedo G, Xiao H, van der Knaap E. 2013. Genome-wide identification, phylogeny and expression analysis of SUN, OFP and YABBY gene family in tomato. Molecular Genetics and Genomics, 288. 10.1007/s00438-013-0733-0.

44. Jung KH, Ko HJ, Nguyen MX, Kim S-R, Ronald P, An G. 2012. Genome-wide identification and analysis of early heat stress responsive genes in rice. Journal of Plant Biology 55, 458–468. 10.1007/s12374-012-0271-z.

45. Klap C, Yeshayahou E, Bolger AM, Arazi T, Gupta SK, Shabtai S, Usadel B, Salts Y, Barg R. 2017. Tomato facultative parthenocarpy results from SlAGAMOUS-LIKE 6 loss of function. Plant Biotechnology Journal 15, 634–647. 10.1111/pbi.12662.

46. Kover PX, Valdar W, Trakalo J, Scarcelli N, Ehrenreich IM, Purugganan MD, Durrant C, Mott R. 2009. A Multiparent Advanced Generation Inter-Cross to fine-map quantitative traits in Arabidopsis thaliana. PLoS Genetics 5, e1000551. doi: 10.1371/journal.pgen.1000551.

47. Kotak S, Larkindale J, Lee U, von Koskull-Döring P, Vierling E, Scharf KD. 2007. Complexity of the heat stress response in plants. Current Opinions in Plant Biology 10, 310–316. doi: 10.1016/j.pbi.2007.04.011.

48. Larkindale J, Hall JD, Knight MR, Vierling E. 2005. Heat stress phenotypes of Arabidopsis mutants implicate multiple signaling pathways in the acquisition of thermotolerance. Plant Physiology 138, 882–897. doi: 10.1104/pp.105.062257.

49. Levy A, Rabinowitch HD, Kedar N. 1978. Morphological and physiological characters affecting flower drop and fruit set of tomatoes at high temperatures. Euphytica 27, 211–218.

50. Li W, Chetelat RT. 2010. A pollen factor linking inter- and intraspecific pollen rejection in tomato. Science 330, 1827–1830. doi: 10.1126/science.1197908.

51. Li J, Zhang H, Hu J, Liu J, Liu K. 2012. A heat shock protein gene, CsHsp45.9, involved in the response to diverse stresses in cucumber. Biochemical Genetics 50, 565–78. doi: 10.1007/s10528-012-9501-9.

52. Lin Y, Laosatit K, Chen J, Yuan X, Wu R, Amkul K, Chen X, Somta P. 2020. Mapping and Functional Characterization of Stigma Exposed 1, a DUF1005 Gene Controlling Petal and Stigma Cells in Mungbean (*Vigna radiata*). Frontiers in Plant Sciences 11, 575922. doi: 10.3389/fpls.2020.575922.

53. Liu YS, Zamir D. 1999. Second generation *L. pennellii* introgression lines and the concept of bin mapping. Report of the Tomato Genetics Cooperative 49, 26–30.

54. Lu Y, Liu L, Wang X, Han Z, Ouyang B, Zhang J, Li H. 2015. Genome-wide identification and expression analysis of the expansin gene family in tomato. Molecular Genetics Genomics 291, 597–608. doi: 10.1007/s00438-015-1133-4.

55. Marathi B, Jena KK. 2015. Floral traits to enhance outcrossing for higher hybrid seed production in rice: present status and future prospects. Euphytica 201, 1–14.

56. Mata-Nicolás E, Montero-Pau J, Gimeno-Paez E, et al. 2020. Exploiting the diversity of tomato: the development of a phenotypically and genetically detailed germplasm collection. Horticultural Research 7, 66. 10.1038/s41438-020-0291-7.

57. Mazzucato A, Papa R, Bitocchi E, et al. 2008. Genetic diversity, structure and marker-trait associations in a collection of Italian tomato (*Solanum lycopersicum* L.) landraces. Theoretical and Applied Genetics, 116, 657–669. 10.1007/s00122-007-0699-6.

58. Meng S, Xiang H, Yang X, et al. 2023. Analysis of YUC and TAA/TAR gene families in tomato. Horticulturae, 9, 665. 10.3390/horticulturae9060665.

59. Muqaddasi QH, Lohwasser U, Nagel M, Börner A, Pillen K, Röder MS. 2016. Genome-Wide Association Mapping of Anther Extrusion in Hexaploid Spring Wheat. PLoS ONE 11, e0155494. 10.1371/journal.pone.0155494.

60. Nnungu SI, Uguru MI. 2014. Expression of heterosis in floral traits and fruit size in tomato (*Solanum lycopersicum*) hybrids. Journal of Tropical Agriculture, Food, Environment and Extension 13, 24 – 29. doi: 10.4314/as.v13i3.4.

61. Opedal ØH, Hildesheim LS, Armbruster WS. 2022. Evolvability and constraint in the evolution of three-dimensional flower morphology. American Journal of Botany 109, 1906–1917. doi:10.1002/ajb2.

62. Pan C, Ye L, Zheng Y, Wang Y, Yang D, Liu X, Chen L, Zhang Y, Fei Z, Lu G. 2017. Identification and expression profiling of microRNAs involved in the stigma exsertion under high-temperature stress in tomato. BMC genomics 18, 843. doi: 10.1186/s12864-017-4238-9.

63. Pan C, Yang D, Zhao X, Jiao C, Yan Y, Lamin-Samu AT, Wang Q, Xu X, Fei Z, Lu G. 2019. Tomato stigma exsertion induced by high temperature is associated with the jasmonate signalling pathway. Plant Cell Environment 42, 1205–1221. doi: 10.1111/pce.13444.

64. Pascual L, Desplat N, Huang BE, Desgroux A, Bruguier L, Bouchet JP, Le QH, Chauchard B, Verschave P, Causse M. 2015. Potential of a tomato MAGIC population to decipher the genetic control of quantitative traits and detect causal variants in the resequencing era. Plant Biotechnology Journal 13, 565–77. doi: 10.1111/pbi.12282.

65. Peralta IE, Spooner DM. 2000. Classification of wild tomatoes: a review. Kurtziana 28, 45–54.

66. Pons C, Casals J, Palombieri, S, et al. 2022. Atlas of phenotypic, genotypic and geographical diversity present in the European traditional tomato. Horticulture Research, uhac112. 10.1093/hr/uhac112.

67. Pons C, Casals J, Brower M, et al. 2023. Diversity and genetic architecture of agro-morphological traits in a Core Collection of European traditional tomato. Journal of Experimental Botany 74, 5896–5916. doi: 10.1093/jxb/erad306.

68. Purcell S, Neale B, Todd-Brown K, et al. 2007. PLINK: a tool set for whole-genome association and population-based linkage analyses. The American Journal of Human Genetics 81, 559–575. doi: 10.1086/519795.

69. Ranc N, Muños S, Xu J, Le Paslier MC, Chauveau A, Bounon R, Rolland S, Bouchet JP, Brunel D, Causse M. 2012. Genome-wide association mapping in tomato (*Solanum lycopersicum*) is possible using genome admixture of *Solanum lycopersicum* var. *cerasiforme*. G3 (Bethesda) 2, 853–64. doi: 10.1534/g3.112.002667.

70. Riccini A, Picarella ME, De Angelis F, Mazzucato A. 2021. Bulk RNA-Seq analysis to dissect the regulation of stigma position in tomato. Plant Molecular Biology 105, 263–285. 10.1007/s11103-020-01086-9.

71. Rick CM. 1983. Transgression for exserted stigma in a cross with *L. pimpinellifolium*. Tomato Genetics Cooperative Reports 33, 13.

72. Rodríguez GR, Muños S, Anderson C, Sim SC, Michel A, Causse M, Gardener BB, Francis D, van der Knaap. 2011. Distribution of SUN, OVATE, LC, and FAS in the tomato germplasm and the relationship to fruit shape diversity. Plant Physiology 156, 275–285. doi: 10.1104/pp.110.167577.

73. Sacco A, Ruggieri V, Parisi M, Festa G, Rigano MM, Picarella ME, Mazzucato A, Barone A. 2015. Exploring a Tomato Landraces Collection for Fruit-Related Traits by the Aid of a High-Throughput Genomic Platform. PLoS One 10, e0137139. doi: 10.1371/journal.pone.0137139.

74. Saeed A, Hayat K, Khan AA, Iqbal S. 2007. Heat Tolerance Studies in Tomato (*Lycopersicon esculentum* Mill.). International Journal of Agriculture & Biology 9, 649–652.

75. SAS Institute. 1988. SAS users guide: statistics. Cary, NC, USA: SAS Institute.

76. Sato S, Peet MM, Thomas JF. 2000. Physiological factors limit fruit set of tomato (*Lycopersicon esculentum* Mill.) under chronic, mild heat stress. Plant, Cell & Environment 23, 719–726.

77. Scott JW, George WL. 1980. Breeding and combining ability of heterostylous genotypes for hybrid seed production in *Lycopersicon esculentum* Mill. Euphytica 29, 135–144. 10.1007/BF00037259

78. Semel Y, Nissenbaum J, Menda N, Zinder M, Krieger U, Issman N, Pleban T, Lippman Z, Gur A, Zamir D. 2006. Overdominant quantitative trait loci for yield and fitness in tomato. Proceedings National Academy of Sciences USA 103, 12981–6. doi: 10.1073/pnas.0604635103.

79. Shang L, Song J, Yu H, et al. 2021.A mutation in a C2H2-type zinc finger transcription factor contributed to the transition toward self-pollination in cultivated tomato. Plant Cell 33, 3293–3308. doi: 10.1093/plcell/koab201.

80. Sun H, Fan HJ, Ling HQ. 2015. Genome-wide identification and characterization of the *bHLH* gene family in tomato. BMC Genomics, 16, 9. 10.1186/s12864-014-1209-2.

81. Tam V, Patel N, Turcotte M, Bossé Y, Paré G, Meyre D. 2019. Benefits and limitations of genome-wide association studies. Nature Reviews Genetics 20, 467–484. 10.1038/s41576-019-0127-1.

82. Turchi L, Baima S, Morelli G, Ruberti I. 2015. Interplay of HD-Zip II and III transcription factors in auxin-regulated plant development, Journal of Experimental Botany 66. 5043–5053. 10.1093/jxb/erv174.

83. van der Knaap E, Chakrabarti M, Chu YH, et al. 2014. What lies beyond the eye: the molecular mechanisms regulating tomato fruit weight and shape. Frontiers in Plant Science 5, 227. doi: 10.3389/fpls.2014.00227.

84. Wu S, Xiao H, Cabrera A, Meulia T, van der Knaap E. 2011. SUN regulates vegetative and reproductive organ shape by changing cell division patterns. Plant Physiology 157, 1175–1186. doi: 10.1104/pp.111.181065.

85. Xu J, Ranc N, Muños S, Rolland S, Bouchet JP, Desplat N, Le Paslier MC, Liang Y, Brunel D, Causse. 2013. Phenotypic diversity and association mapping for fruit quality traits in cultivated tomato and related species. Theoretical and Applied Genetics 126. 567–581. doi: 10.1007/s00122-012-2002-8.

86. Xu C, Liberatore KL, Macalister CA, et al. 2015. A cascade of arabinosyltransferases controls shoot meristem size in tomato. Nature Genetics 47, 784–792. doi: 10.1038/ng.3309.

87. Xu J, Driedonks N, Rutten MJ, Vriezen WH, de Boer GJ, Rieu I. 2017. Mapping quantitative trait loci for heat tolerance of reproductive traits in tomato (*Solanum lycopersicum*). Molecular breeding, 37, 58. doi: 10.1007/s11032-017-0664-2.

88. Zhang W, Zhou RG, Gao YJ, Zheng SZ, Xu P, Zhang SQ, Sun DY. 2009. Molecular and genetic evidence for the key role of *AtCaM3* in heat-shock signal transduction in Arabidopsis. Plant Physiology 149, 1773–1784. doi: 10.1104/pp.108.133744.

